# ZIC3 controls the transition from naïve to primed pluripotency

**DOI:** 10.1101/435131

**Authors:** Shen-Hsi Yang, Munazah Andrabi, Rebecca Biss, Syed Murtuza Baker, Mudassar Iqbal, Andrew D. Sharrocks

## Abstract

Embryonic stem cells (ESCs) are pluripotent in nature, meaning that they have the capacity to differentiate into any cell in the body. However, to do so they must transition through a series of intermediate cell states before becoming terminally differentiated. A lot is known about how ESCs maintain their pluripotent state but comparatively less about how they exit this state and begin the transition towards differentiated cells. Here we investigated the earliest events in this transition by determining the changes in the open chromatin landscape as naïve mouse ESCs transition to epiblast-like cells (EpiLCs). Motif enrichment analysis of the newly opening regions coupled with expression analysis identified ZIC3 as a potential regulator of this cell fate transition. Chromatin binding and genome-wide transcriptional profiling confirmed ZIC3 as an important regulatory transcription factor and among its targets are genes encoding a number of transcription factors. Among these is GRHL2 which acts through enhancer switching to maintain the expression of a subset of genes from the ESC state. Our data therefore place ZIC3 at the top of a cascade of transcriptional regulators and provide an important advance in our understanding of the regulatory factors governing the earliest steps in ESC differentiation.

**Highlights:**

- The transcription factor ZIC3 drives gene expression changes in the ESC to EpiLC transition.
- Extensive changes occur in the open chromatin landscape as ESCs progress to EpiLCs.
- ZIC3 activates the expression of a network of transcription factors.
- ZIC3 activated genes in EpiLCs are upregulated in the post-implantation epiblast.

## Introduction

Early embryonic development involves the transition of pluripotent embryonic stem cells through intermediate cell states into the cell lineages that initiate subsequent development events. Using defined *in vitro* conditions, several different states have been identified for mouse ESCs, starting from the naïve ground state, progressing through EpiLCs to establish an epiblast stem cell (EpiSC) state (Hayashi et al., 2011; reviewed in Kalkan and Smith, 2012). Subsequently, EpiSCs can differentiate to the three germ layers, mesoderm, ectoderm and endoderm. Mouse ESCs can be maintained in the naïve ground state in defined media which includes two kinase inhibitors (known as ‘2i’) to block the MEK/ERK and GSK3 signalling pathways (Ying et al., 2008; reviewed in Wray et al., 2010). Withdrawal of 2i, allows the cells to progress to either EpiLCs or EpiSCs by altering culture conditions (Betschinger et al., 2013; Hayashi et al., 2011). The naïve ESCs are thought to represent a model for the pre-implantation epiblast (E3.5-4.5) whereas EpiLCs or EpiSCs cells are models for the post-implantation epiblast (E5.5) (Kalkan et al., 2017).

As ESCs progress from the naïve ground state, large changes are observed in their chromatin landscapes and underlying gene expression programmes (Marks et al., 2012; Factor et al., 2014; reviewed in Habibi and Stunnenberg, 2017). The pluripotent state is maintained through the action of a core set of transcription factors and chromatin regulators that include the well-studied NANOG, KLF4, SOX2, and OCT4, (reviewed in Young, 2011). However, comparatively less is known about the regulators controlling the transition to EpiLCs and EpiSCs. Recently OTX2 was identified as a key transcription factor driving this transition, partly through cooperative interactions with OCT4/POU5F1 (Acampora et al., 2013; Buecker et al., 2014; Yang et al., 2014). Proteomics analysis also identified ZIC2/3 and OCT6/POU3F1 as interacting proteins for OCT4 specifically in EpiLCs (Buecker et al., 2014), suggesting a potential co-regulatory role for these transcription factors in this context. Additional changes occur during the transition to EpiLCs, and in addition to transcriptional regulators, other proteins have been shown to play an important role during this transition such as the extracellular signalling protein, Cripto which controls metabolic reprogramming (Fiorenzano et al., 2016).

To further our understanding of the regulatory networks controlling the transition from the naïve ESC state to EpiLCs, we examined the chromatin accessibility changes accompanying this early transition in mouse ESCs. We focused on areas of dynamic chromatin opening and through DNA binding motif enrichment and associated gene expression data analysis, we identified the transcription factor ZIC3 as an important regulatory transcription factor in this context. ZIC3 controls the expression of EpiLC marker genes such as *Fgf5* and many of the ZIC3 target genes encode transcriptional regulators such as GRHL2 which has an important role in enhancer formation in the transition to EpiLCs. ZIC3 therefore sits at the top of a cascade of regulators that work together to establish and maintain the EpiLC state.

## Results

### Identification of transcription factors involved in the transition to EpiLCs through open chromatin profiling

Cell state transitions are accompanied by changes to the underlying regulatory chromatin landscape (Stergachis et al., 2013). These changes can then be used to infer the potential roles of upstream transcription factors (Sung et al., 2014). To begin to understand the regulatory events occurring during the conversion of naïve mouse ESCs to EpiLCs, we therefore profiled the accessible chromatin landscape of mouse ESCs as they transition to EpiLCs over a 2 day period (Fig. 1A) using ATAC-seq. Gene expression changes at matched time points were also profiled using single cell (sc) RNA-seq from 816 cells. Open chromatin regions were identified at each time point and the resulting peaks consolidated into a single reference dataset (238,236 peaks in total). These peaks were then partitioned between promoter proximal (−2 kb to +0.5 kb), intragenic and intergenic regions to examine whether genomic location affected the overall changes in chromatin accessibility. We then identified regions that showed differential accessibility between any two conditions, giving 3,041 (promoter), 16,510 (intragenic) and 17,306 (intergenic) differentially accessible regions. These regions were then clustered into four broad patterns based on their chromatin opening dynamics (Fig. 1B; Supplementary Fig. S1A and B); regions that increased accessibility at day 1 and became further accessible at day 2 (cluster 1), regions that decreased accessibility at day 1 and became even more inaccessible at day 2 (cluster 2), or regions that transiently opened or closed at day 1 (cluster 3 and 4).

**Figure 1.**
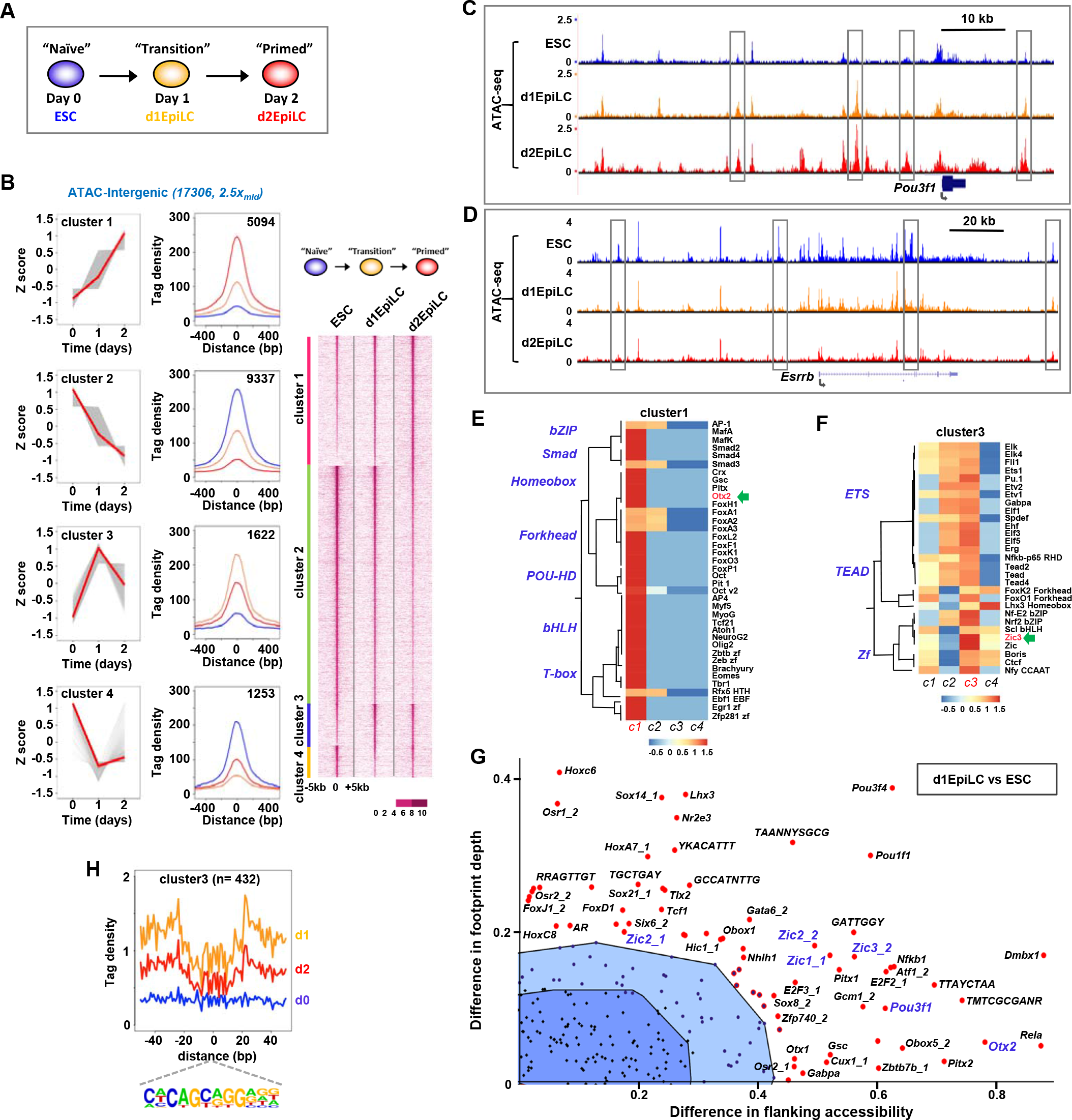
Identification of transcriptional regulators of the ESC to EpiLC transition by open chromatin profiling. (A) Schematic of the experimental time course of the naïve ESC to EpiLC transition. (B) Heat map of the ATAC-seq profiles across a 10 kb window of intergenic regions showing >2.5 fold change in accessibility between any two conditions (right). Average tag densities of each of four identified clusters (middle; blue=ESC, orange=d1EpiLCs, red=d2EpilCs) and average tag density profiles (z scored) are shown across the time course (left). Medians [red] and data for individual peaks [grey] are indicated. (C and D) UCSC genome browser views of the ATAC-seq profiles around the *Pou3f1* and *Esrrb* loci. Dynamically changing peaks are boxed. (E and F) Heat map showing the enrichment of transcription factor binding motifs across each of the open chromatin cluster profiles (z-normalised p-values) for motifs enriched in cluster 1 (E) or cluster 3 (F). (G) BaGFoot analysis of the open chromatin regions in ESCs and d1EpiLCs (using all dynamic intergenic peaks from clusters 1-4 in B). The top right quadrant from the whole plot (see Supplementary Fig. S4) is shown. Motifs showing significant increases in either local accessibility and/or footprint depth are labelled. Non-significant regions are shown in the dark (bag) or light (fence) blue shaded regions. (H) Average tag densities in a 100 bp window surrounding the ZIC3 binding motif (bottom) in ESCs (blue), d1EpiLCs (orange) or d2EpilCs (red) are shown for ATAC-seq peaks from cluster 3.

We next recovered the genes associated with each promoter and matched intergenic peaks to their likely associated genes by using the nearest gene model. Then we compared the changes in expression relative to the changes in open chromatin associated with each gene. By focussing on the differentially changing ATAC-seq peaks and gene expression changes, we observed a good concordance between chromatin opening and gene expression changes (Supplementary Fig. S1C and D). These changes become more marked after 2 days as cells acquire the EpiLC state. The changes we observed in chromatin accessibility profiling therefore generally report on the activity status of associated genes. To further examine whether these accessibility profiles reflected the underlying changes in gene expression, we took the open chromatin peaks in each cluster, and associated them with all genes located at different peak-to-gene distances and calculated the enrichment of resulting set of genes among the equivalent clusters derived from scRNA-seq data. Overall, there was an excellent correlation between the two datasets, with the best matches occurring between similar cluster patterns (Supplementary Fig. S1E and F). Furthermore, each of the ATAC-seq clusters was associated with groups of genes exhibiting a unique set of GO terms (Supplementary Fig. S2A-D). For example, cluster 1 and cluster 2 regions are associated with various developmental terms as might be expected by their sequential changes in the transition to EpiLCs. The regulatory regions of genes encoding transcription factors associated with the two cell states show expected changes during the transition to EpiLCs; several peaks within the *Pou3f1* locus (EpiLC transcription factor) show sequential opening and are found in cluster 1 (Fig. 1C). Conversely, peaks in the *Essrb* locus (ESC transcription factor) shows progressive closing and are found in cluster 2 (Fig. 1D). In contrast, cluster 4 genes are associated with various stem cell processes, consistent with transient regulatory region closing as illustrated by the *Nodal* locus (Supplementary Fig. S2F).

Having established the relevance of open chromatin profiling to gene expression changes, we next wanted to identify the relevant regulators. To that end, we searched the differentially accessible regions for over-represented transcription factor binding motifs. Each of the accessibility clusters has a different repertoire of motifs (Fig. 1E and F; Supplementary Fig. S3A-F) with ZEB1, KLF4, ZIC3 and TCF3 being the most enriched binding motifs for clusters 1-4, respectively. Interestingly, OTX2 binding motifs were identified in cluster 1 regions, which is consistent with the fact that these regions become sequentially more open in the transition to EpiLCs and the known role for OTX2 in driving early ESC fate decisions (Yang et al., 2014; Buecker et al., 2014). We were particularly interested in cluster 3, as these regions are characterised by transient opening at day 1, suggesting an important role in the transition towards EpiLCs. To further interrogate the underlying transcription factor networks in this cluster, we used BaGFoot (Baek *et al.*, 2017) to identify transcription factor motifs that exhibit increased footprint depth (and hence occupancy) and/or local DNA accessibility. Numerous motifs were identified including those for several homeodomain and ZIC proteins (Fig. 1G; Supplementary Fig. S4A). ZIC binding motifs had previously been associated with OCT4/POU5F1 binding regions in EpiLCs (Buecker et al. 2014), therefore we focussed on this binding site, and more closely examined the chromatin accessibility surrounding this motif at the three differentiation time points. Clear footprints were observed in open chromatin clusters 1 and 3 around this motif in day 1 (d1) EpiLCs and the depth and local accessibility mirrored the general accessibility profiles of these clusters across different time points (Fig. 1H; Supplementary Fig. S4B). We also compared the open chromatin of day 2 (d2) EpiLCs to ESCs and applied a similar analysis. Multiple motifs were again identified as becoming more accessible and potentially more occupied including those for OTX2 and the ZIC transcription factors (Supplementary Fig. S5).

Together these results establish the dynamics of chromatin accessibility changes accompanying the transition from ESCs to EpiLCs and identify ZIC transcription factors as likely important players in controlling gene regulation during this transition.

### ZIC3 is transiently upregulated during the transition to EpiLCs

There are multiple members of the ZIC transcription factor family; therefore we determined their relative expression levels during the transition to EpiLCs. *Zic1* and *Zic4* are not expressed to appreciable levels, whereas *Zic2* and *Zic5* show progressively increased expression at day 1 and day 2 of the differentiation timecourse (Fig. 2A). However, *Zic3* shows a transient increase in expression at day 1, which is even more pronounced at the protein level (Fig. 2B). These findings are supported by single cell RNA-seq analysis, where *Zic3* expression is enriched in the d1EpiLCs (Fig. 2C). In contrast, OTX2 expression shows fewer dynamic changes and is increased at day 1 and remains at a stable level in day 2 EpiLCs (Fig. 2A-C). The ZIC transcription factors therefore show dynamic changes in their expression that accompanies the transition to EpiLCs, and the transient expression kinetics of ZIC3 in particular indicates that this is a likely candidate for controlling the transition phase.

**Figure 2.**
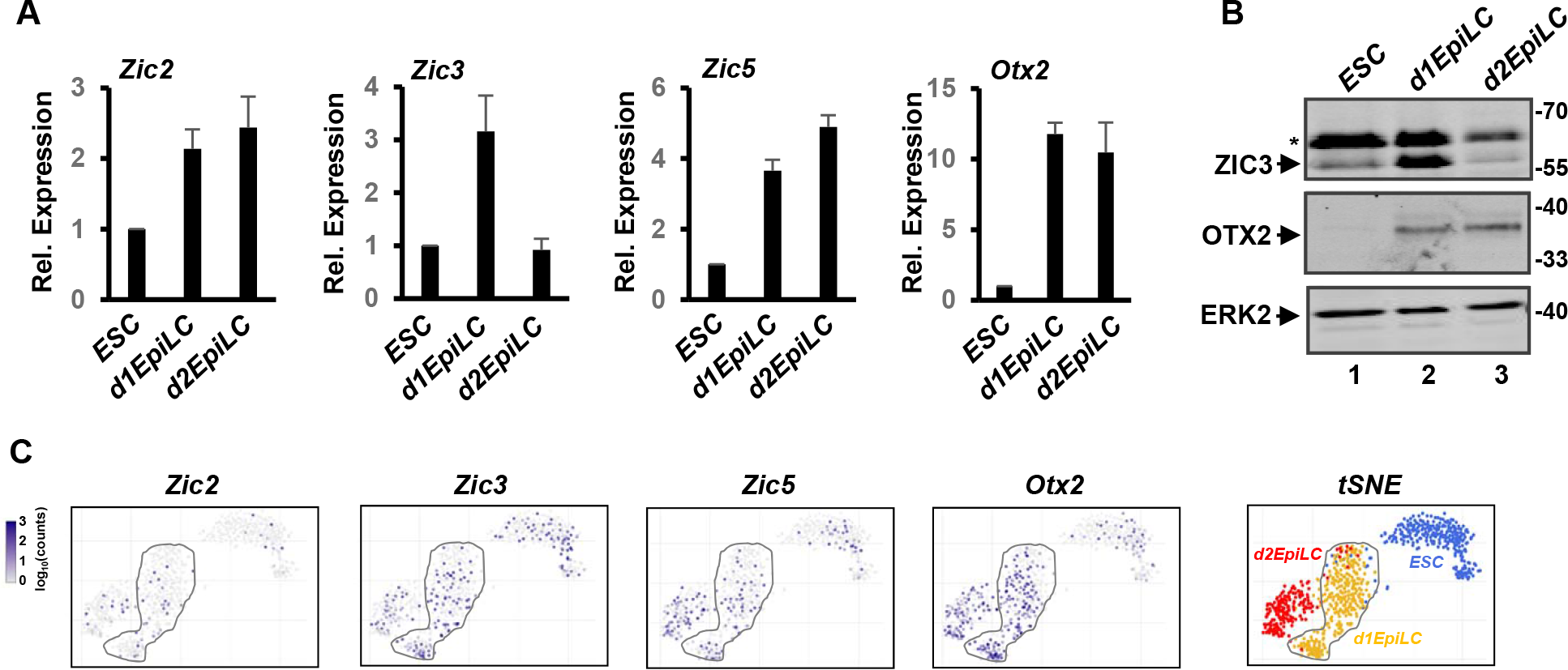
Expression profiles of Zic transcription factors in ESCs and EpiLCs. (A) RT-qPCR analysis of *Zic2*, *Zic3*, *Zic5* and *Otx2* expression in the indicated cell states (n=3). (B) Western blot analysis of ZIC3, OTX2 and ERK2 expression. The asterisk marks a non-specific band. (C) Single cell RNA-seq analysis of *Zic2*, *Zic3*, *Zic5* and *Otx2* expression. t-SNE analysis of the entire scRNA-seq dataset is shown on the right, with the originating cell types colour coded. d1EpiLCs are circled.

### Determination of the ZIC3 cistrome

Next, we focussed on ZIC3 and, as a first step in determining its regulatory potential, we identified its genome-wide binding profile using ChIP-seq. Initially we focussed on the transition state on day 1 and identified 4,724 high confidence ZIC3 bound regions (Fig. 3A; Supplementary Table S1). The majority of these are located in inter- and intra-genic regions and the ZIC3 binding regions are associated with 5,216 target genes based on the nearest neighbour model. Consistent with a role for ZIC3 in cell fate changes, these target genes are enriched in GO terms for many differentiation processes, and signalling pathways such as the BMP and STAT pathways (Fig. 3B). As expected, the ZIC binding motif is highly enriched in these regions along with more moderate enrichment for several other transcription factors, including ESSRB and SOX proteins, which have previously been implicated in regulatory activities in stem cells (Fig. 3C).

**Figure 3.**
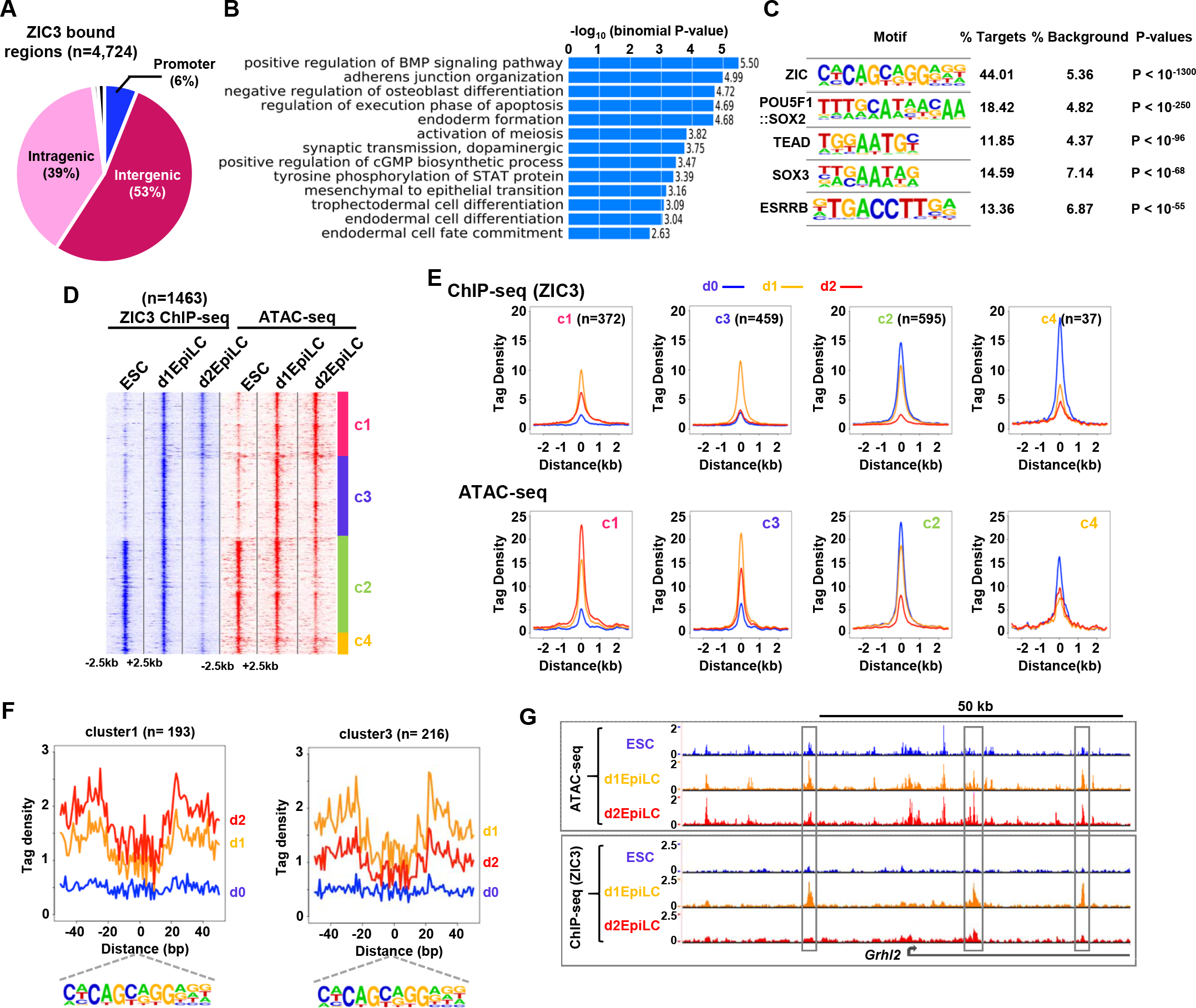
ChIP-seq analysis of ZIC3 genomic binding. (A) Genome-wide distribution of ZIC3 binding sites in d1EpiLCs. Promoter is defined as −2.5 to + 0.5 kb. (B) Gene ontology analysis of ZIC3-associated genes (biological process). (C) Top 5 enriched motifs found in the ZIC3 binding regions. (D) Heat map of the ZIC3 ChIP-seq profiles across a 5 kb window of all inducible ATAC-seq peaks (>2.5 fold change for intra-/inter-genic peaks and >2 fold change for promoter peaks) (left). The corresponding ATAC-seq signals at each ZIC3 binding region is shown on the right. Data are clustered (clusters c1-c4) according to ATAC-seq signals. (E) Average tag densities of ZIC3 binding peaks from each of four identified clusters in each cell population (blue=ESC, orange=d1EpiLCs, red=d2EpilCs) for ZIC3 ChIP-seq signal (top) or ATAC-seq signal (bottom). (F) Average ATAC-seq tag densities in an 80 bp window surrounding the ZIC3 motif (bottom) in cluster c1 (left), or cluster c3 (right). Data from ESCs (blue), d1EpiLCs (orange) or d2EpilCs (red) are shown. (G) UCSC genome browser views of the ATAC-seq (top) and ZIC3 ChIP-seq (bottom) profiles around the *Grhl2* locus. Dynamically changing ZIC3 binding peaks are boxed.

To uncover ZIC3 binding dynamics and link these to the changing chromatin accessibility profiles, we performed additional ChIP-seq experiments for ZIC3 in ESCs and day 2 EpiLCs. Replicate experiments showed good concordance (Supplementary Fig. S6A) and clustered together in PCA analysis (Supplementary Fig. S6B). Overall, the binding dynamics showed transiently increased occupancy of ZIC3 at day 1 which was reduced back to a lower level, at day 2 (Supplementary Fig. S6C and D). These occupancy changes were accompanied by a transient increase in chromatin opening across the binding regions at day 1 (Supplementary Fig. S6C and D). This transient opening could be observed in more detail when analysing the cut frequencies around the ZIC3 binding motifs (Supplementary Fig. S6E). This was particularly marked when considering the ZIC3 binding regions, which were not already occupied in ESCs (d1 unique peaks; Supplementary Fig. S6E, right). To gain further insight into the relationship between binding and chromatin accessibility dynamics, we focussed on the chromatin regions showing changes in accessibility between any two cell conditions (Fig. 1B; Supplementary Fig. S1A and B). ZIC3 binding across these regions generally mirrors the changes in chromatin accessibility (Fig. 3D and E). For example, in cluster c3, ZIC3 binding is strongly enhanced at day 1 as chromatin accessibility increases, and is lost again at day 2 as chromatin accessibility decreases again. However, in cluster c1, ZIC3 binding becomes reduced at day 2 (consistent with its decreased expression) but chromatin accessibility increases, indicating a disconnection between ZIC3 binding kinetics and chromatin accessibility in these regions. This may reflect other factors acting to maintain or enhance the chromatin accessibility status at these regions. The changes in chromatin accessibility were also revealed by focussing on the cleavage events around the ZIC3 motifs located in the ZIC binding regions (Fig. 3F; Supplementary Fig. S6F and G). When considering all ZIC3 binding regions both the depth and local accessibility is transiently enhanced in day 1 EpiLCs (Supplementary Fig. S6F). By focussing on different subclusters, different patterns could be discerned (Fig. 3F Supplementary Fig. S6G). For example, footprint depth and local accessibility in regions belonging to cluster 3 show highest levels in day 1 EpiLCs as expected from the increased ChIP-seq signals in these regions. This behaviour is exemplified by the *Grhl2* locus where several ZIC3 peaks are maximally present in d1EpiLCs, and this transient increase is accompanied by chromatin opening at the same loci (Fig. 3G; Supplementary Fig. S6H).

Collectively these data reveal a dynamically changing ZIC3 cistrome during the transition from ESCs to EpiLCs. These dynamic changes are accompanied by underlying changes to the open chromatin landscape surrounding their sites.

### ZIC3-dependent gene regulatory events

Having established the ZIC3 cistrome and its dynamic changes, we next asked whether ZIC3 influences gene expression through these binding events. We depleted *Zic3* (Supplementary Fig. S7 A and B) and determined the changes in transcriptome at the day 1 EpiLC transition state. 452 genes changed expression (>1.2 fold), with 53% showing reduced expression following *Zic3* depletion, which is consistent with a potential activator role for ZIC3 (Fig. 4A; Supplementary table S2). GO term analysis of the ZIC3 regulated genes, revealed enrichment of categories including cell adhesion alongside several developmental terms, various signalling pathways and “regulation of transcription” (Fig. 4B; Supplementary Fig. S7C). Indeed, among these genes, there are numerous transcription factors and signalling pathway components (Fig. 4A) indicating large changes in the regulatory systems in the cells. Two additional notable examples are the genes encoding the cell surface proteins, EPCAM and PECAM1, which show reciprocal changes in expression following *Zic3* depletion (Fig. 4A, highlighted in purple) and are usually expressed at distinct times during the differentiation process, with PECAM1 being an ESC marker and EPCAM an EpiLC marker (Fig. 4C and D). We next examined the expression of the ZIC3 activated genes across single cells that had been ordered by pseudo time analysis (Trapnell *et al*. 2014) of scRNAseq data. Each cell was scored for expression of each target gene in a binary manner and the overall fraction of genes expressed per cell determined. There is a clear increase in expression of the ZIC3 regulon as cells progress towards d1EpiLCs and beyond (Fig. 4E). However, this is not apparent in cells that do not co-express *Zic3* (Fig. 4F). This association is further reflected by the increases in co-expression levels of the ZIC3 activated genes in d1 and d2EpliCs, which is not observed in cells with low *Zic3* expression (Fig. 4G).

**Figure 4.**
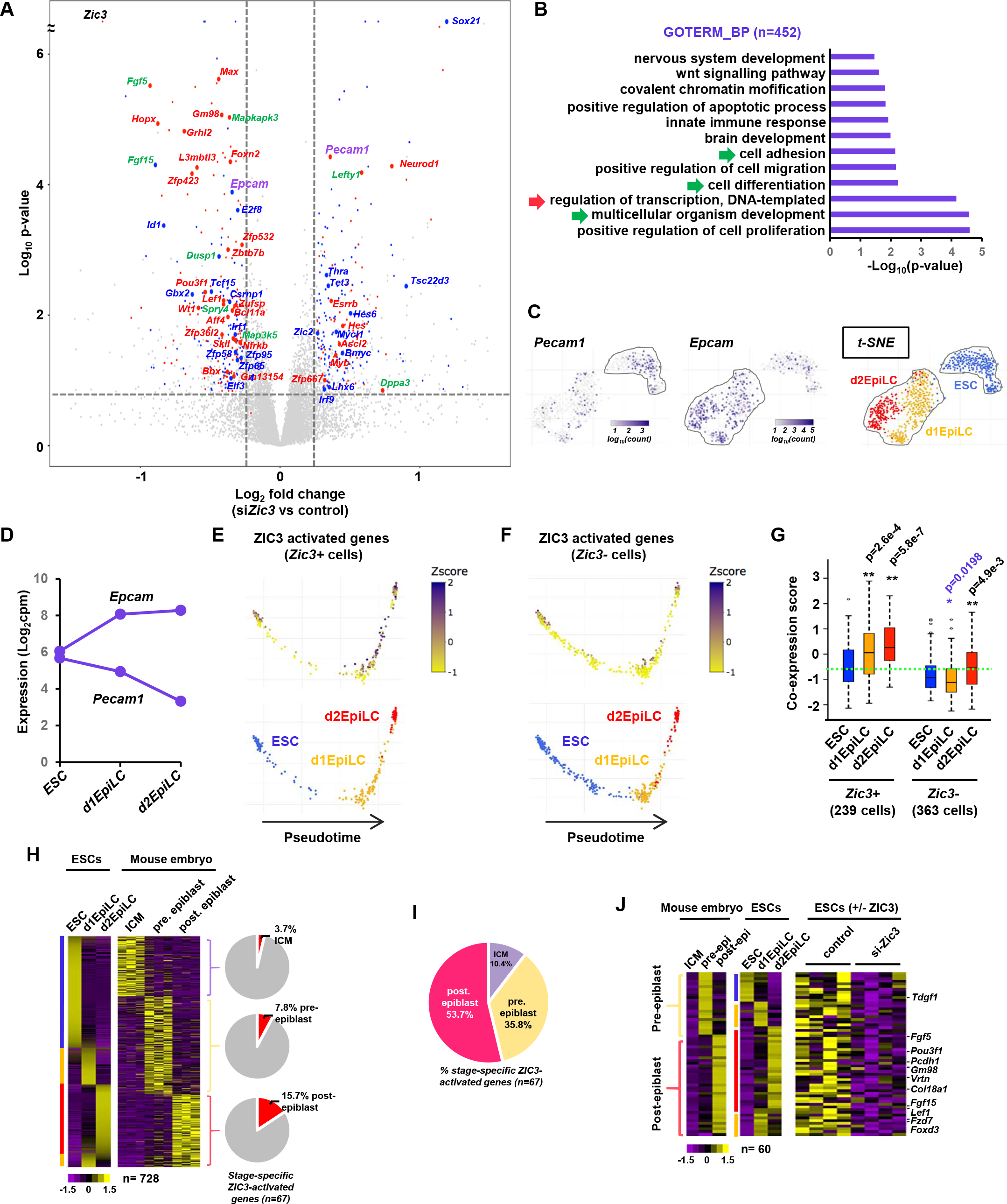
Identification of Zic3-regulated genes. (A) Volcano plot showing changes in gene expression in d1EpiLCs following *Zic3* knockdown by siRNA. Gene names are colour coded for transcription factors (red for direct and blue for indirect targets) and signaling molecules (green). *Pecam1* and *Epcam* are shown in purple. (B) Gene ontology analysis of all ZIC3 regulated genes for the biological process (BP) category. (C) Expression of *Pecam1* and *Epcam* at single cell level. Data are plotted as log_10_ counts per cell, and superimposed on t-SNE analysis of the entire RNA-seq dataset for three time points (right). (D) Expression of *Epcam* and *Pecam1* from aggregated single cell RNA-seq analysis in the indicated cell populations and shown as log_2_ counts per million base pairs (cpm). (E and F) Pseudotime analysis of ESCs, d1EpiLCs and d2EpilCs based on the entire scRNA-seq dataset (bottom). The binarised expression (transformed z-score) of the ZIC3 activated genes (n=240) is plotted on top of these profiles (top) in cells that either show *Zic3* expression (E) or lack the expression of *Zic3* and *Otx2* (F). Pseudotime analysis was initially performed with all cells but in each case only cells exhibiting the *Zic3* expression characteristics are shown. (G) Boxplot showing the co-expression scores for the ZIC3 activated genes in ESCs, d1EPiLCs and d2EpilCs. Cells are split according to whether *Zic3* is expressed or not. Horizontal lines represent median score, and dotted green line is the median score in ESCs. (H) Heatmaps showing the expression levels of genes categorised as uniquely expressed in ICM, pre-epiblast or post-epiblast (right; Boroviak et al., 2015) and the corresponding expression levels in the aggregated single cell RNA-seq from ESCs, d1EPiLCs and d2EpilCs (left). The heatmap is sorted based on the scRNAseq data from the ESC-derived cells at each of the expression clusters from mouse embryos. Data are row z-normalized for each dataset. The pie charts show the proportions of each of the stage-specific gene sets that are activated by ZIC3. (I) Pie chart showing the proportions of ZIC3-activated lineage-specific genes from each stage of embryonic development. (J) Heatmap showing the effect of *Zic3* depletion on the pre-and post-epiblast stage-specific genes in d1EpiLCs (right) and the heatmaps for the corresponding gene expression levels in early embryonic developmental stages (left) or ESCs, d1EpiLCs and d2EpilCs (centre). Genes shown are from the red quadrants of the bottom two pie charts in part H. All heatmaps are individually z-normalised.

Finally, we asked whether the ZIC3-regulated genes are relevant in the context of early mouse embryonic development. We analysed the clusters of genes that exhibit peak expression levels at each stage of embryonic development (Boroviak et al. 2015) and first compared the data to our own RNA-seq data from aggregated single cell analysis of ESCs, d1EpiLCs and d2EpiLCs. Overall, there is good concordance between the datasets with ESCs being most similar to the ICM, d2EpiLCs resembling the post-implantation epiblast and d1EpiLCs representing an intermediate state (Fig. 4H). Next, we asked whether ZIC3 is involved in regulating the expression of any of these genes that act as markers of early embryonic development. Importantly, when we superimposed our *Zic3* depletion dataset on top of these clusters, there was a sequential increase in the number of ZIC3-activated genes among the marker genes expressed maximally at each stage of embryonic development (Fig. 4H, right; Fig. 4I). Among these genes are known markers and regulators of differentiation in the pre-epiblast and post-epiblast stages such as *Foxd3* and *Fgf5* (Fig. 4J)(Hanna et al., 2002; Khoa et al., 2016). Thus ZIC3 activates the expression of a large number of marker genes that are characteristic of the mouse pre- and post-implantation epiblast.

Next, we identified the direct target genes for ZIC3. To achieve this, and gain further insight into the likely direct roles of ZIC3, we took our ChIP-seq data and associated ZIC3 ChIP peaks with nearby genes. By intersecting this with our RNA-seq data we uncovered a total of 207 directly regulated target genes for ZIC3 (Supplementary table S2). The majority of these are activated by ZIC3 (65%) (Supplementary Fig. S7D) suggesting a role for ZIC3 in upregulating gene expression during the transition to EpiLCs. Indeed, the directly activated ZIC3 target genes show an overall increase in expression during the transition to EpiLCs and this is particularly marked in d1EpiLCs, with over 70% of these genes showing upregulation (Fig. 5A, left). One notable directly activated target gene is *Wt1*, which encodes a bifunctional splicing factor and sequence-specific transcription factor which is known to function post-transcriptionally to regulate developmental RNAs in mouse ESCs (Bharathavikru et al., 2017) (Fig. 5B). Reciprocally, we observe the opposite for the “directly repressed” genes, albeit to a lower level of significance (Fig. 5A, right), leaving open the possibility that ZIC3 may be a bifunctional transcription factor. We also examined co-expression of ZIC3 directly activated genes using AUCell analysis, which is specifically designed to identify co-expression across single cells (Aibar et al., 2017). More frequent expression of the ZIC3 regulon was observed in the d1EpiLCs (Fig. 5D), consistent with a role for ZIC3 in controlling gene expression at this transition point. Finally, we asked whether the ZIC3 regulon has predictive potential in determining cell types from scRNA-seq data and showed that the 135 directly activated ZIC3 target genes are not only able to separate ESCs from EpiLCs (Fig. 5E) but are also are able segregate cells from different embryonic stages using scRNA-seq data derived from mouse embryos (Mohammed et al., 2017), with E4.5 epiblast cells forming a distinct compact cluster (Supplementary Fig. S7E).

**Figure 5.**
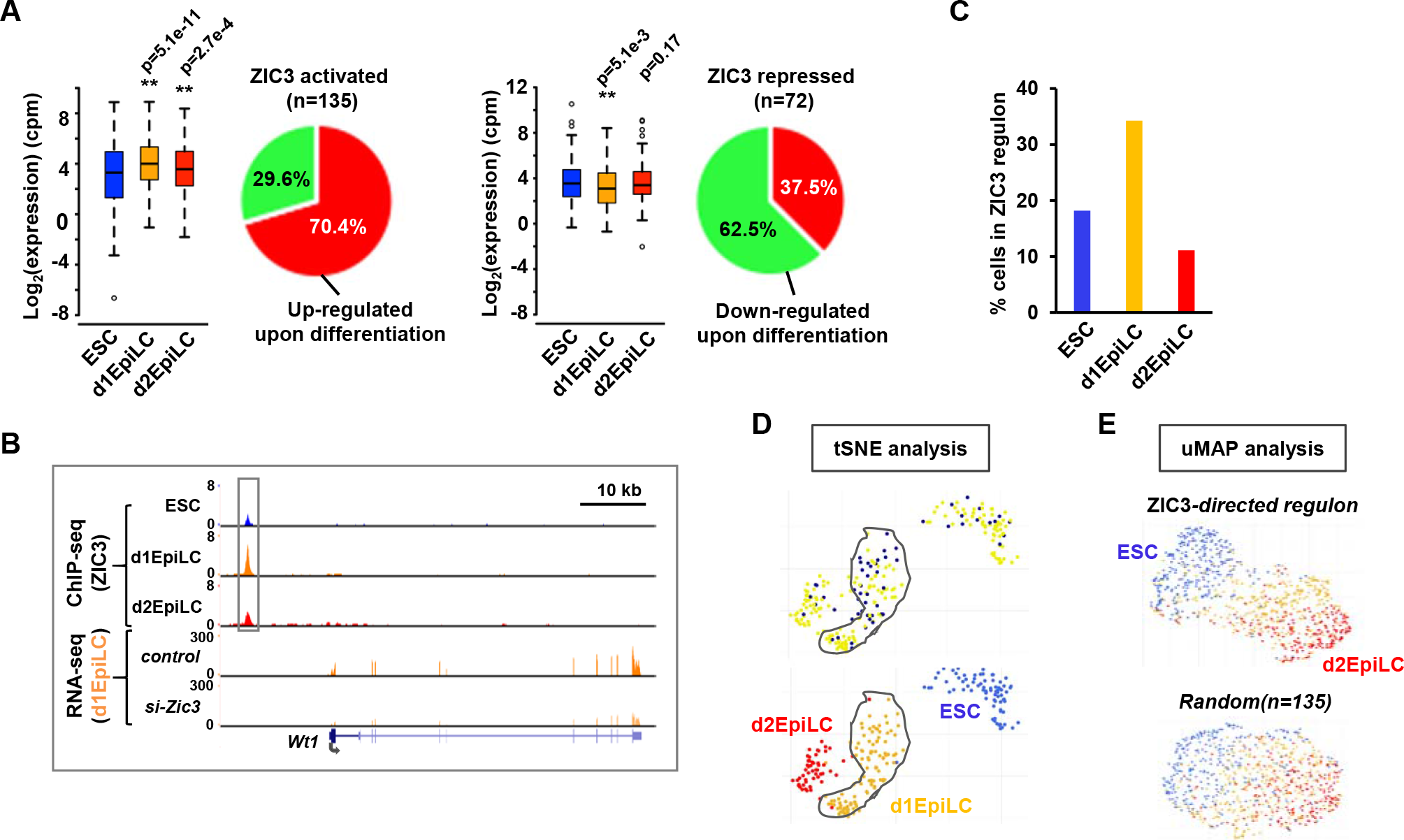
The direct ZIC3 target gene network. (A) Boxplots of the expression of directly regulated ZIC3 target genes (ie bound by ZIC3) in ESCs, d1EpiLCs and d2EpilCs for activated (top) or repressed (bottom) genes. The proportions of direct ZIC3 target genes increasing and decreasing expression in d1EpiLCs upon differentiation from ESCs are shown in the pie charts on the right. (B) UCSC genome browser views of the ZIC3 ChIP-seq (top) and RNA-seq (bottom) profiles around the *Wt1* locus. The major ZIC3 binding peak is boxed. (C) AUCell analysis of the expression of the directly activated ZIC3 target gene regulon in ESCs, d1EPiLCs and d2EpilCs. The percentage of cells is shown from each stage of differentiation that exhibit co-expression of the ZIC3 regulon. (D) Single cell RNA-seq analysis of the ZIC3 regulon expression. Data are mapped (blue marked cells; top) on top of tSNE analysis of the entire RNA-seq dataset (bottom), with the originating cell types colour coded. d1EpiLCs are circled. Only *Zic3* positive cells are shown. (E) uMAP analysis of the single cell RNA-seq ESCs, d1EPiLCs and d2EpilCs using either the 135 ZIC3-activated direct target genes in the ZIC3 regulon (top) or 135 randomly selected genes (bottom) to drive the clustering. Cells are colour coded according to their known origins (blue=ESCs, orange=d1EpiLCs, red=d2EpiLCs).

ZIC3 is therefore involved in directly controlling the expression of a set of target genes that are generally upregulated in the transition from ESCs to EpiLCs and in the pre- and post-epiblast stages in the developing embryo.

### ZIC3 triggers a complex downstream transcriptional regulatory network

To further understand the mechanisms through which ZIC3 affects downstream transcriptional programmes that result in the EpiLC phenotype, we examined the functions of several of its target genes. Many of the ZIC3-regulated genes encode transcription factors, suggesting that ZIC3 acts mechanistically to trigger subsequent waves of changes in the transcriptome, mediated by these intermediary transcription factors. A large number of these are direct targets including *Wt1*, *Lef1*, *Grhl2* and *Pou3f1*. Further analysis of the single cell RNA-seq data demonstrates that a subset of these transcription factors is strongly co-expressed in d1EpiLCs (Fig. 6A and Supplementary Fig. S8A and B). This transcription factor network is largely absent in ESCs but maintained in d2EpiLCs. To begin to understand the impact of these transcription factors on downstream gene expression patterns, we asked whether we could find evidence for their DNA binding motifs in open chromatin regions associated with genes that show elevated expression in d2EpiLCs. We focussed on regions that show increases in accessibility during the transition to EpiLCs (ie clusters 1 and 4 in Fig. 1B) and found that a number of motifs are over-represented in these regions, including those for ZIC3 and OTX2 (Fig. 6B). Importantly, motifs are also over-represented for a number of transcription factors encoded by ZIC3-activated genes, such as TCF15, WT1, LEF1, GRHL2 and POU3F1 (Fig. 6B) consistent with a potential role for these transcription factors in enhancing the expression of these genes in EpiLCs. As an alternative approach, we used BaGFoot on all of the open regions that are associated with the genes showing enhanced expression in the transition from d1EpiLCs to d2EpiLCs and found that GRHL2 and POU3F1 binding motifs were among the motifs showing strong evidence for increased footprint depth and localised chromatin opening in EpiLCs (Fig. 6C). Given the strong presence of binding motifs in the regulatory regions of potential target genes, we next sought evidence for regulatory activity of the corresponding transcription factors. We focussed on GRHL2 as this has recently been shown to play an important role in switching enhancer usage during the transition of ESCs to EpiLCs (Chen et al., 2018). GRHL2 is encoded by a direct ZIC3 target gene (see Fig. 3G), suggesting a potential functional hierarchy with ZIC3 acting upstream from GRHL2 in a transcriptional cascade. This hierarchy predicts that ZIC3 depletion should have a similar effect on downstream gene expression profiles as depletion of GRHL2. We therefore focussed on a set of directly activated GRHL2 target genes (ie bound by GRHL2) which showed the largest decreases in expression in EpiLCs following loss of GRHL2 expression (Chen et al., 2018). Importantly, the majority of these were downregulated upon depletion of ZIC3 in d1EPiLCs and/or d2EpiLCs (Fig. 6D), consistent with a transcription factor relay network whereby ZIC3 controls GRHL2 expression to subsequently influence downstream gene expression. More generally, ZIC3 controls the expression of a set of transcription factors that are able to generate a cascade effect on gene expression at later stages of ESC differentiation.

**Figure 6.**
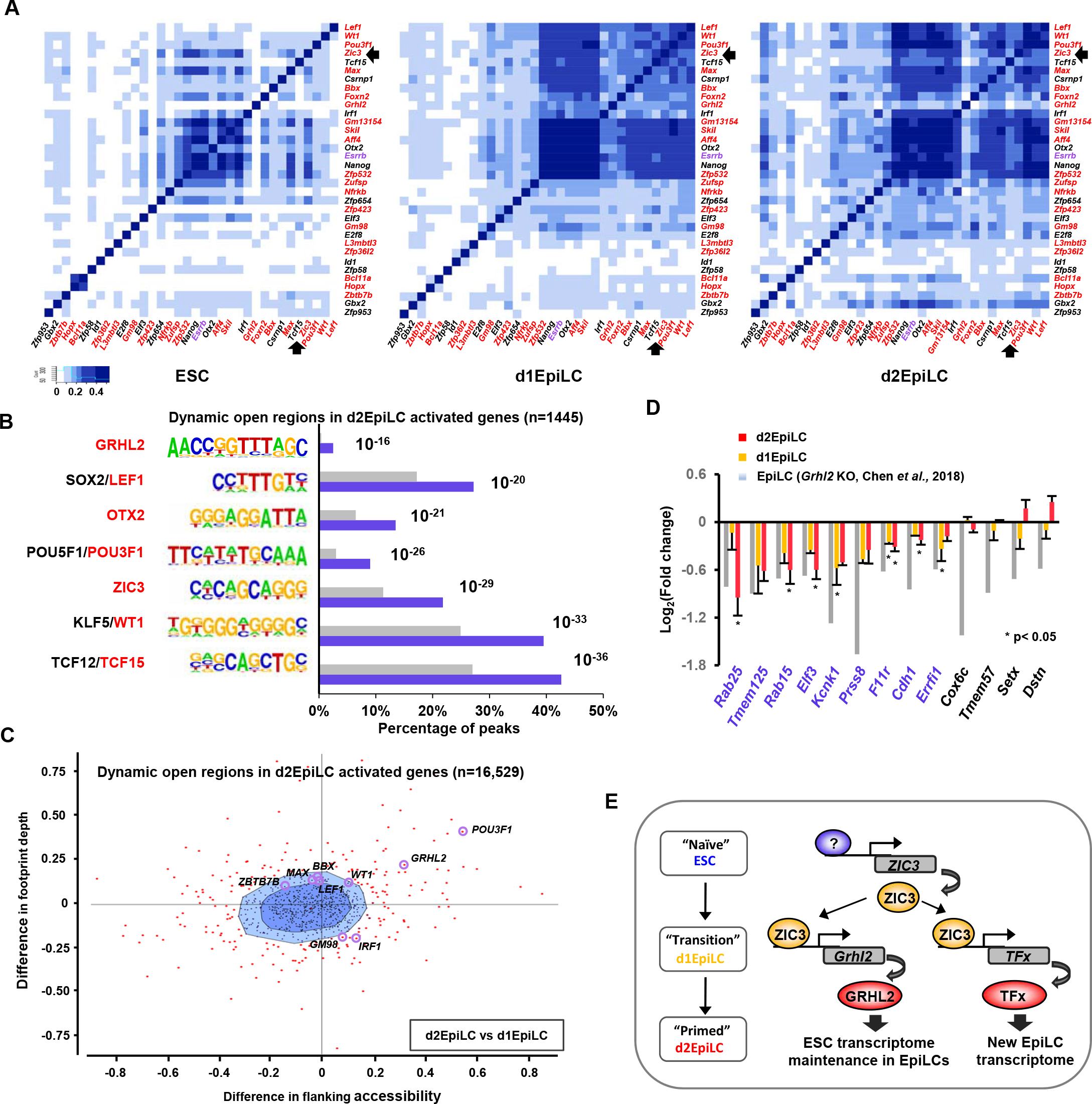
ZIC3 regulated transcription factors control downstream gene expression programmes. (A) Jaccard’s similarity plots of the co-expression of the indicated transcription factor-encoding ZIC3 activated genes in ESCs (left), d1EPiLCs (middle) and d2EpilCs (right). Direct ZIC3 targets are shown in red and the location of *Zic3* is highlighted with an arrow. (B) Enrichment of DNA motifs within open chromatin regions associated with genes that change expression >2 fold from d1EpiLCs to d2EpiLCs. Only dynamically opening inter- and intra-genic ATAC-seq peaks (ie from clusters 1 and 4, Fig. 1B and Supplementary Fig. S1B) were analysed (n=1,445). The percentage of each motif present in each set of peaks (blue) and the genomic background (grey) is indicated, and the p-value is shown next to each of the columns. (C) BaGFoot analysis of all of the open chromatin regions in d2EpiLCs which are associated (125 kb to +125 kb; n=16,529) with genes which increase in expression (>2 fold) in going from d1EpiLCs to d2EpiLCs. Motifs corresponding to binding sites for transcription factors encoded by ZIC3-activated genes are labelled. (D) The expression of the indicated direct GRHL2 target genes (Chen et al., 2018) following depletion of *Grhl2* (grey bars) or *Zic3* in d1EpiLCs (orange bars) and d2EpiLCs (red bars). Data are shown based on the fold changes seen in RNA-seq data. Asterisks show significantly changing expression levels (p-value <0.05) and standard deviations are indicated (n≥3). (E) Model showing the transcriptional events centred on ZIC3 during the transition from ESCs to EpiLCs. GRHL2 maintains the ESC transcriptome (Chen et al., 2018) whereas other ZIC3-regulated transcription factors likely contribute to the newly established EpiLC transcriptome.

## Discussion

The transcription factor networks controlling maintenance of the pluripotent state in ESCs are relatively well understood. However, it is less clear how naïve ESCs begin differentiation by transitioning through the EpiLC state. The transcription factor OTX2 was previously shown to control enhancer activation during the transition from naïve ESCs towards EpiLCs (Acampora et al., 2013; Yang et al., 2014; Buecker et al., 2014). Here we took an unbiased approach using ATAC-seq to uncover novel transcriptional regulators of this transition. We focused on ZIC3 which exhibits transient expression kinetics and chromatin binding as cells change fate to EpiLCs. ZIC3 plays a key role in controlling gene expression during differentiation to EpiLCs, and in particular a large number of genes encoding signaling molecules and transcription factors. Through activating the expression of transcription factor encoding genes, ZIC3 acts at a pivotal point in a transcriptional cascade which determines the EpiLC phenotype (Fig. 6E). For example, the ZIC3 regulated transcription factor, GRHL2 has been shown to play an important role in maintaining a stem cell-specific gene expression programme as cells progress to the EpiLC state through an enhancer switching mechanism (Chen et al., 2018). Other ZIC3 regulated transcription factors play a role in controlling other gene expression programmes during differentiation such as TCF15 which has previously been shown to be important in priming EpiSCs for differentiation (Davies et al., 2013).

Previous studies suggested that ZIC transcription factors are likely involved in early developmental decisions in naïve ESCs. ZIC2 and 3 were identified as interactors of OCT4 in EpiLCs, suggesting a role in cooperative transcriptional regulation as the OCT4 cistrome is remodeled in the transition from ESCs (Buecker et al., 2014). Indeed, we provide further support for this model as we identified an enrichment of OCT4-like binding motifs in ZIC3 binding regions (Fig. 3C). However, other motifs are enriched in the ZIC3 bound regions, suggesting a broader cooperativity with a wide range of transcription factors. Interestingly, ZIC3 has also been implicated in the maintenance of pluripotency in ESCs (Lim et al., 2007). However in the latter study, the ESCs were maintained in the presence of serum and LIF, conditions which mimic the EpiSC state rather than naïve ESCs. Our data are therefore generally consistent with a role for ZIC3 in ESCs, but point to ZIC3 acting at an early stage during the transition from naïve ESCs. ZIC3 knockout mice exhibit early embryonic developmental defects prior to gastrulation, leading to defects in left-right patterning (Ware et al., 2006). Moreover, mutation of ZIC3 in humans causes a syndrome known as X-linked heterotaxy, where similar patterning defects are observed (reviewed in Bellchambers and Ware, 2018). It is possible that these developmental defects arise due to the changes we observe at the earliest cell fate transitions from naïve ESCs. Indeed, consistent with this early role, many of the ZIC3 regulated target genes show peak expression during the transition from the ICM through to the pre-and post-implantation epiblast in the embryo (see Fig. 4H).

ZIC transcription factors can bind similar DNA binding motifs (Badis et al., 2009; reviewed in Hatayama and Aruga, 2018) and hence there is the potential for further cross talk between these factors at the level of chromatin binding. As ZIC3 exhibits transient activation kinetics, and ZIC2/5 expression is maintained at later stages, it is possible that some of the functions of ZIC3 are maintained and/or expanded by these factors as EpiLCs differentiate further. ZIC2 was previously shown to act in concert with the Mbd3/NuRD complex in ESCs to cause transcriptional repression and loss of ZIC2 affected subsequent ESC differentiation (Luo et al., 2015). However, the ESCs in this study were cultured under conditions that create the EpiSC state. Further work in this cell type also implicated ZIC2 as an important player in the transcriptional regulatory circuits in these cells (Matsuda et al., 2017). Moreover ZIC1 and ZIC2 have been shown to play a role much later in development in the context of the neuronal gene expression programme in cerebellar granule neurones (Frank et al., 2015). Future studies will be needed to unravel whether ZIC3 function in the early ESC transitions are modified by other ZIC family members later in the differentiation and development programme.

It is unclear whether ZIC transcription factors are transcriptional activator or repressor proteins. Work on ZIC2 suggests a repressive role (Luo at el. 2015). In contrast, the majority of the directly regulated ZIC3 target genes (65%) are activated by ZIC3, although a substantive proportion are repressed. It is possible that like many transcription factors, ZIC3 can adopt different roles at different regulatory regions, and in this context may poise genes in the transition state for subsequent activation in EpiLCs. Nevertheless, through its transcriptional regulatory activities, ZIC3 plays an important role in controlling the transition from naïve ESCs to the more advanced EpiLC state. Through activating genes encoding transcription factors such as GRHL2 it contributes to maintaining a plastic state which retains stem-cell properties but is poised for subsequent differentiation. It is highly likely that other ZIC3 regulated transcription factors play equally important roles in creating this flexible regulatory environment.

## Materials and Methods

### Tissue culture, RNA interference and RT-PCR

Mouse *Rex1GFP*d2 ES cells were maintained as described previously in NDiff 227 media (Takara Bio Europe SAS, Y40002) containing 2i inhibitors (CHIR99021 and PD0325901; Miltenyi Biotec, 130-106-539 and 130-106-549) on dishes coated with gelatin (Millipore, ES-006-B) (Yang *et al.*, 2012). The d1EpiLCs were created by plating 2.5×10^4^ cells/cm^2^ on dishes coated with bovine plasma fibronectin (5 μg/ml; Sigma, F1141) and then growing for 1 day in NDiff N2B27 media containing bFGF (12ng/ml; R&D system, 233-FB-025), activin A (20 ng/ml; Peprotech,120-14E-10), and KnockOut Serum Replacement (1%; ThermoFisher, 10828010) (Hayashi *et al.*, 2011). For d2EpiLCs, the half volume of medium was removed and replenished with freshly prepared medium, and growth continued for a further day. RNAi was performed as described previously (Yang *et al.*, 2012).

Real time RT-qPCR was carried out as described previously (O’Donnell *et al.* 2008). Data were normalised for the geometric mean expression of the control genes *hmbs* and *ppia*. The primer-pairs used for RT-PCR are listed in Supplementary Table S3.

### Western blot analysis

Western blotting was carried out with the primary antibodies; Erk2 (137F5; Cell Signalling, 4695), Otx2 (ProteinTech., 13497-1-AP) and ZIC3 (Abcam, ab222124). All experiments were carried out in 12-well plates. The lysates were directly harvested in 2xSDS sample buffer (100 mM Tris.Cl pH 6.8, 4% SDS, 20% glycerol, 200 mM DTT and 0.2% bromophenol blue) followed by sonication (Bioruptor, Diagenode). The proteins were detected using a LI-COR Odyssey Infrared Imager as described previously (Yang *et al.*, 2012).

### ATAC-seq assays

The cells were dissociated from the plates with Accutase (Sigma, A6964) for 3 minutes at 37°C. ATAC-seq samples and libraries were generated as described previously (Buenrostro *et al.*, 2015) except the nuclei were prepared using 100 μl of ice cold Nuclei EZ lysis buffer (Sigma, N3408). The nuclei pellets were resuspended in 10 μl H_2_O and nuclei were counted. The tagmentation reaction was performed with 50 thousand nuclei and 2.5 μl of Tn5 transposase (0.5 μM) in 25 μl reaction volumes for 30 mins at 750 rpm at 37°C. The tagmented genomic DNA was purified by using miniElute Reaction Cleanup kits (Qiagen, 28204) and eluted in 10.5 μl. The libraries were generated by 9 cycles of PCR reaction using adaptor primers (Nextera Index kit; Illumina, FC-121-1012) and NEBNext high fidelity 2x PCR master mix (NEB, M0541), followed by two-sided size selection by Ampure XP beads purification (0.4x reaction volume then 1.2x reaction volume; Beckman Coulter Agencourt, A63881). The typical yield is between 150-300 ng. The sequencing was performed on an Illumina Next-seq genome analyser according to the manufacturer’s protocols. ATAC-seq data have been deposited in the ArrayExpress repository (http://www.ebi.ac.uk/arrayexpress/) under accession number: E-MTAB-7207.

### ChIPmentation assays

For ChIP-seq using the ChIPmentation method (Schmidl *et al.* 2015), the cells (1.4-2 × 10^7^ cells sufficient for 5 ChIPmentation experiments) were dissociated with Accutase (Sigma, A6964) for 3 mins at 37°C and fixed in 1% formaldehyde in 0.03%BSA/F12 for 10 min at room temperature. After quenching with 0.125 M glycine, cells were pelleted and the pellets were washed with 0.03%BSA/PBS. The nuclei were lysed in lysis buffer (10 mM Tris-Cl pH 8.0, 10 mM NaCl, 0.2% NP40 and 1 tablet of Complete protease inhibitor cocktail (Thermo Scientific) per 50 ml for 20 mins at 4°C. The nuclei were counted, snap frozen in liquid N_2_ and stored at −80°C. Prior to ChIPmentaion, nuclei pellets were resuspended in H_2_O and topped up with 0.25% SDS (25×10^6^ cells/ml; 130 μl/aliquot). The nuclei solution was then sonicated 3 times, 10 cycles (30s on/off) at 4°C (Bioruptor, Diagenode). The IP solution was prepared through sequential dilution by adding 1.5 volumes of equilibration buffer (10 mM Tris.Cl pH8.0, 140 mM NaCl, 0.6 mM EDTA, 1% triton X-100, 0.1% Na-deoxycholate and 0.1% SDS) and 0.92 volumes of TopUp buffer (10 mM Tris-Cl pH8.0, 140 mM NaCl, 1 mM EDTA, 1% triton X-100, 0.1% Na-deoxycholate and 0.05% SDS).

ChIPmentation assays were performed essentially as described previously (Schmidl *et al.* 2015). Briefly, 100 μl of ZIC3 antibody solution (1 μg; Abcam, ab222124) was cross-linked to the protein A beads were incubated with 100 μl IP solution (1.2×10^6^ nuclei) at 4°C overnight. The beads were sequentially washed twice with 250 μl of low salt buffer, high salt buffer, LiCl wash buffer and once in 150 μl of 10mM Tris-Cl pH 8.0. Next, the tagmentation reactions (25 μl) were performed with 1 μl of Nextera Tn5 transposase (Nextera kit; Illumina, FC-121-1030) in tagmentation buffer (33 mM Tris-OAc pH 7.8, 66 mM potassium-OAc, 10 mM Mg-OAc and 16% DMF) at 1200 rpm at 37°C for 10 mins. Ice cold low salt buffer (150 μl) was immediately added to stop the enzymatic reaction on ice for 5 mins. The beads were then washed twice with 150 μl of low salt buffer and TE. The tagmented ChIPed samples were then resuspended in 48 μl of ChIPmentation elution buffer (10 mM Tris.Cl pH8.0, 300 mM NaCl, 5 mM EDTA and 0.4% SDS) and incubated with 2 μl of proteinase K (20 mg/ml; Ambion, AM2546) at 55°C for 1 hr and then at 65°C overnight. The beads were further incubated for 1hr at 55°C with proteinase K (1 μl 20 μg/ml)/ChIPmentation elution buffer (19 μl). The combined eluates were topped up with 300 μl of ERC and ChIPed DNA purified by a mini Elute kit (Qiagen). ChIPed DNA was eluted in 10.5 μl of EB buffer. The sequencing libraries were generated by 12 cycles of PCR reaction using adaptor primers (Nextera Index kit; Illumina, FC-121-1012) and NEBNext high fidelity 2x PCR master mix (NEB, M0541), followed by two-sided size selection by Ampure XP beads purification (0.65x reaction volume then 1.2x reaction volume; Beckman Coulter Agencourt, A63881). The typical yield is between 150-300 ng. The sequencing was performed on an Illumina Hi-seq 4000 genome analyser according to the manufacturer’s protocols. ZIC3 ChIPmentation-seq has been deposited in the ArrayExpress repository (http://www.ebi.ac.uk/arrayexpress/) under accession number: E-MTAB-7208.

### RNA-seq assays

Total RNA was prepared using RNAeasy Plus Mini kit (RNase-free DNase set; Qiagen, 74134) according to the manufacturer’s protocols except extra “in column” DNase digestion was performed (Qiagen, 79254). Libraries for RNA-seq were generated using the Illumina TruSeq RNA library prep kit v2 (Illumina, RS-122-2001) and sequencing was performed on an Illumina Hi-seq 4000 genome analyser according to the manufacturer’s protocols (ArrayExpress accession: E-MTAB-7206).

### Single cell (sc) RNA-seq assays

Single cell RNA-seq was performed on the ICELL8 single-cell RNA-seq system as described previously (Goldstein *et al.*, 2017) except that cryogenically frozen cells were used (ArrayExpress accession: E-MTAB-7211).

### Bioinformatics and statistical analysis

All software was run with default settings, unless otherwise indicated. Raw sequencing reads (76-nt length; paired end) were trimmed and filtered using Trimmomatic v0.32 with paired-end mode to remove adapters, truncated reads (3’) and reads with <25 nucleotides (TRAILING:5 SLIDINGWINDOW:4:15 MINLEN:25; Bolger *et al.*, 2014). Filtered reads were mapped against National Center for Biotechnology Information build 37/mm9 of mouse genome using Bowtie2 v2.3.0 (allow up to two mismatches, *-X 2000 and ȁdovetail*; Langmead *et al.* 2009). Unmapped pairs (-F 4) were discarded using SAMtools v1.3.1 (Li *et al.*, 2009). Reads were then de-duplicated using the *MarkDuplicates* function of the Picard tools (http://broadinstitute.github.io/picard/). Only reads that were uniquely mapped to the genome were preserved (MAPQ ≥30). The reads mapped to the mitochondrial genome (sed ‘/chrM/d’) and overlapping with mm9 blacklist regions (intersectBed -v) were removed. The normalised tag density profiles were generated using HOMER (annotatePeak.pl; Heinz *et al.*, 2010) and were plotted using a customised R script. Heatmaps were generated using Java treeview (Eisen *et al.* 1998). The UCSC tracks were generated by genomeCoverageBed for ATAC-seq normalised to total reads in peaks (RIPs) (BEDtools; Quinlan & Hall, 2010), MACS2 for ChIPmentation normalised to total tags (Zhang *et al.*, 2008) or RSeQC for RNA-seq (Wang *et al.*, 2012).

### ATAC-seq data analysis

ATAC-seq peaks (open-chromatin regions) were called using MACS2 (Zhang *et al.*, 2008) on individual replicates with the following parameters: -q 0.01 --nomodel -- shift -75 --extsize 150. The high confidence peak sets were selected from biological replicates using the *intersectBed* function from BEDTools (Quinlan & Hall, 2010) with parameters −f 0.50, −r. This ensures a reciprocal overlap of >50% between the two peaks being selected. To get a union set of peaks from all three conditions (ESC, d1EpiLC and d2EpiLC), high confidence MACS peaks from each condition were merged using *mergePeaks* module from HOMER (d=100; Heinz *et al.*, 2010) so only a single peak was retained when two or more peaks from different conditions had peak to peak distance <100 bp). All downstream analysis was based on this union set of 238,236 peaks.

For identifying differentially accessible regions and fuzzy cMeans clustering, the union set of peaks was divided into promoter (−2 kb to +0.5 kb), intragenic-(defined by peaks located within mm9 protein coding regions) and intergenic-regions (all remaining peaks). Read counts for all peaks in the union set were obtained using the *annotatePeaks* module of HOMER package (Heinz *et al.*, 2010) and were quantified using edgeR (Robinson *et al.*, 2010). Fuzzy cMeans clustering using the R Mfuzz package (Kumar *et al.*, 2007) was then performed on each set of ATAC-accessible peaks identified in the Promoter, Intergenic and Intragenic regions, respectively. Initially, the Fuzzy cMeans clustering was performed to classify peaks into 12 clusters, which were subsequently merged into 4 clusters upon manual inspection. The final differentially accessible peaks were filtered based on EdgeR analysis (minimal CPM ≥ 4 in any of the three conditions), q < 0.05 and fold change ≥ 2 (promoter peaks) ≥ 2.5 (intergenic- and intragenic-peaks) (on at least one pairwise comparison between conditions).

To determine motif enrichment in clustered regions, over represented transcription factor motifs in each of the four clusters were identified using findMotifsGenome module of the HOMER package (Heinz *et al.*, 2010). Motifs were then clustered using Fuzzy cMeans clustering and were also assigned to their respective families using the STAMP tool (Mahony *et al.*, 2007). The relative enrichment scores of the clustered motifs were then transformed to Z-scores and plotted as heatmaps using the R *pheatmap* package.

To identify the transcription factors undergoing substantial changes in occupancy levels and chromatin accessibility around their binding sites between the transition states we used BaGFoot (Baek *et al.*, 2017) software on the clustered ATAC-seq peaks. ATAC-seq peaks from all 4 clusters were merged for a reliable detection of footprint depth with robust statistical significance. We collected transcription factors from the JASPAR (Khan *et al.*, 2018) database, which were manually curated to exclude transcription factors from non-vertebrate species, giving us 872 transcription factor motifs. The mouse genome (mm9) was scanned for motif occurrences of these transcription factors using Find Individual Motif Occurrences (*FIMO*) (Grant *et al.*, 2011) as recommended by the software (1.5 M motif threshold count). We performed pairwise comparisons for the transition states (d1EpiLC vs ESC and d2EpiLC vs ESC) and calculated the changes in accessibility and footprint-depth. Results are displayed as bagplots.

### ChIPmentation data analysis

ChIPmentation data was compared to input chromatin and peaks were called on each replicate using MACS2 v2.1.1 using parameters: *--keep-dup all -q 0.01 -g mm -f BAMPE -B --SPMR --call-summits* (Zhang *et al.*, 2008). The high confident peak set (peaks identified in both biological replicates) was selected using *mergePeaks* module from HOMER (d=400, peak summit distance=400; Heinz *et al.*, 2010). Similarly, the *mergePeaks* (d=250, peak summit distance=250) was used to subset peaks that overlapped with differential accessible ATAC-peaks.

Motif discovery and the significance of discovered motifs was performed by HOMER (findMotifsGenome.pl; Heinz *et al.*, 2010) using the sequences within ± 100 bp around the binding region summits, using the default background setting i.e., sequences randomly selected from the genome with the same GC content as the target sequences.

Nearest genes were assigned to peaks and the Gene Ontologies (GO) were analysed using GREAT (McLean *et al.* 2010). Genomic distributions were determined using HOMER (Heinz *et al.*, 2010).

### RNA-seq data analysis

A manually curated gtf file was built for expression quantification of all datasets. Briefly, the gtf (vM1) file for mm9 from the GENCODE website was downloaded and genes specified by transcript_type (protein_coding, lincRNA and antisense) were retained. In addition, genes missing from the GENCODE gtf file but in ENSEMBL gtf file were added to our list. After manual filtering and inspection, the gtf file comprises of 25,875 unique ensembl id’s and 25,753 unique gene symbols.

Filtered paired-end reads were mapped to the mouse genome (mm9 assembly) using STAR v2.5.3a (Dobin et al., 2013) with the manually curated mm9-gtf file and default parameters. Ribosomal RNA (rRNA) reads were removed from the mapped files. Read counts for each sample were quantified using HTSeq v0.9.1 (Anders et al., 2015), which estimates number of reads mapped to each gene. The raw read counts from the HTSeq were subsequently used to quantify the differential expression levels for genes using DESeq2 v1.18.1 (Anders et al., 2010). Data were taken as significant if a fold change of >1.2 and p-value <0.05 was obtained. Additional genes were included in our analysis above this p-value threshold if they changed expression in a consistent direction in paired samples and also exhibited a mean fold change >1.2. For volcano plots, log_2_ fold changes (FC) of differentially expressed genes were plotted against their log_10_ p-values using the inbuilt function of the R statistical package. The Gene Ontology (GO) analyses were performed using DAVID (Huang *et al.*, 2009).

### Single cell transcriptomics

Single cells from three samples (ECS, d1EpiLCs and d2EpiLCs) were captured and isolated using the ICELL8 single cell system. A custom script was used to perform assignment and error correction of cell barcodes/UMIs, low quality reads trimming and to run a cross species contamination checking. After the QCs, reads were aligned to a customized mouse reference genome of mm9 using STAR aligner (v2.4.2a). Reads aligning to genes were counted using HTSeq (v0.6.1.p1) with setting the stranded option to “yes”. This count matrix was then used for the downstream analysis of the dataset using statistical computing programming language R.

We implemented two measures of cell quality control (cell QC) based on library size and number of expressed genes. If the total read count of a cell is below 3× median absolute deviation (MAD) of the dataset then the cell was filtered out. Similarly, if the total number of genes expressed by a cell is lower than 3x MAD those cells were also filtered out. A further QC was done to filter out any cells that are outliers in terms of library size on the higher end as this could indicate doublets of cells. After all these filtering steps, a total of 816 cells (from 869) were left for downstream analysis.

Subsequently, the lowly expressed genes were filtered out if their average counts are less than 0.05 (raw counts) meaning a gene has to be expressed in at least 5% of cells with 1 read count or higher counts in smaller number of cells still accounting for 5% equivalent cells, which gave us 12,695 genes for downstream analysis. The normalised count data is represented as counts per million where the size factors are used to calculate the effective library size. These size factors were defined from the actual library size after centering to unity. We used the calculateCPM function from the scater package (McCarthy *et al.*, 2017) to perform this normalisation.

To identify the highly variable genes (HVGs) we first estimated the total variance in expression of each gene which is then decomposed into technical and biological components. We fitted a mean-variance trend to the expression of endogenous genes and then took those genes that have a larger biological variance component with an FDR value less than 0.05 as our HVGs. 283 genes were identified as HVGs for this dataset. These HVGs were then used to construct dimensionality reduction processes using PCA. For the t-SNE plot (van der Maaten *et al.*, 2008), 10 principal components from this PCA are given as input and the perplexity is set to 60. In addition, the theta (a parameter for speed/accuracy tradeoff) is set to 0.01 to increase the accuracy of the plot. For generating the Uniform Manifold Approximation and Projection (UMAP) plots (McInnes *et al.*, 2018), 10 PCs are taken as input.

We used the monocole (v2) package to perform the pseudotime estimation (Trapnell *et al*. 2014). As we know the three stages of cells in our samples we use this information to identify the order of the genes. Genes that are differentially expressed between ESCs, d1EpiLCs and d2EpiLCs with a q-value less than 7.5e-08 were identified and subsequently used to order cells. We then apply DDRTree method to reduce the dimension of the dataset (Qiu *et al.*, 2017). The pseudotime trajectory is visualized in the reduced dimension.

Co-expression scores across single cell RNA-seq data were calculated by first giving a binary score to the expression of each gene in each cell. These binary scores were summed for each cell, and then z transformed. The data are shown as box plots for ESCs, d1EpiLCs and d2EpiLCs in Fig. 4G after correcting for the numbers of cells at each condition.

To examine the Jaccard Similarity Index (JSI), Jaccard’s distance was computed for the binary co-expression matrix of all ZIC3 activated TFs in each cell type (ESC, d1EpiLC and d2EpiLC2) using the R package *ade4* (Dray and Dufour, 2007). The dissimilarity matrix was then converted to a similarity matrix by using the expression JSI= 1-(JD)^2^ (where JD= Jaccard’s distance) and the JSI based data were plotted as heatmaps. First, the d1EpiLC data was clustered on both row and columns using hierarchical clustering using heatmap.2 from R gplots package. The heatmaps generated for ESC and d2EpiLC data were plotted using the same gene ordering that was obtained from clustering of the d1EpiLC data.

The Matthew’s Correlation Coefficient (MCC) for the co-expression data of ZIC3 activated genes that encode transcription factors, were computed using a custom R script. The correlation matrix for d1EpiLC was then ordered for the first principal component using R package *corrplot* (Wei and Simko, 2017) and plotted as a heatmap after clustering the rows and columns. As in the JSI plots, the gene order from the d1EpiLC heatmap was retained and the heatmaps for ESC and d2EpiLC data were plotted without further clustering.

Beeswarm plots for Jaccard Similarity Indices of Zic3 activated target genes in each cell type were plotted using R package *Beeswarm* (Elkund, 2016; https://rdrr.io/cran/beeswarm).

To interrogate the co-expression of genes in the ZIC3 regulon (ZIC3 targets) in our scRNA-seq data, the AUCell module from SCENIC R package was used (Aibar *et al.*, 2017). AUCell calculates the enrichment of gene-sets (regulon) as area under the recovery curve (AUC) based on the rankings of all genes expressed in a particular cell. The AUC threshold was then determined and subsequently used to mark whether the cells contained an active- or inactive-regulon. This binary data was then visualised by superimposing onto t-SNE plots.

To verify the importance of the ZIC3 regulon during mouse embryogenesis, a scRNA-seq dataset generated during early mouse gastrulation was used (Mohammed *et al.*, 2017). The raw reads were mapped to the same custom mm9 gft file and analysed as described above. HVGs were used to generate the PCAs and 14 PCs were used as input for the t-SNE plot. For the t-SNE plot, the perplexity was set to 60 and theta to 0.01. In addition, 135 direct ZIC3-activated genes were selected as input to cluster cells using the t-SNE plot with the same perplexity and theta value. As a control, 135 randomly selected genes were used.

To generate pseudo-bulk RNA datasets from single cell data, the aggregated counts of each gene from each cell of the scRNA-seq were generated and quantified using the edgeR (Robinson *et al.*, 2010). The Fuzzy cMeans clustering was performed to generate 4 broad expression clusters as described above. 8659 genes were selected for further downstream analysis (CMP ≥ 2, fold change > 1.5 at any pairwise comparison).

To correlate Fuzzy cMeans-generated clusters of ATAC-seq and RNA-seq data (supplemental figure S1E and F), we first identified genes whose TSSs lie within a given genomic distance constraint from any peak within a given cluster of ATAC-seq peaks (A_i_). Next we take the RNA-seq based gene clusters (R_j_), and calculate the intersection of R_j_ with A_i_ (observed set). The expected set of genes was defined as all genes within a given genomic distance of an ATAC-seq cluster from a randomly selected number (according to comparator expression cluster size) of genes extracted from the background population (all mm9 genes). The p-values for the enrichment were subsequently calculated using a hypergeometric test between the observed and expected datasets. This calculation is repeated for all to all combinations between ATAC-seq peak clusters and gene expression clusters, and for each of pre-defined set of genomic distances relative to the TSS (8 bins within the range of +/−10 kb to +/−250 kb from the centre of the peak). The resulting p-values are log-transformed (−log10(p-values)) and shown as heatmaps in figures.

## Acknowledgements

We thank Karren Palmer and Mairi Challinor for excellent technical assistance; staff in the Genomic Technologies, and Bioinformatics facilities; Catherine Millar, Hilary Ashe and members of our laboratories for comments on the manuscript and stimulating discussions. This work was funded by the BBSRC and Wellcome Trust (ADS and SM-B) and the MRC grants MR/M008908/1 and MR/M012174/1 (MI).

